# Predicting individual learning trajectories in zebrafish via the free-energy principle

**DOI:** 10.1101/2025.08.06.668947

**Authors:** Takuya Isomura, Yuki Tanimoto, Makio Torigoe, Hitoshi Okamoto, Hideaki Shimazaki

**Affiliations:** Brain Intelligence Theory Unit, RIKEN Center for Brain Science, 2-1 Hirosawa, Wako, Saitama 351-0198, Japan; RIKEN Center for Brain Science, 2-1 Hirosawa, Wako, Saitama 351-0198, Japan; Faculty of Science and Engineering, Waseda University, 2-2 Wakamatsu-cho, Shinjuku-ku, Tokyo 162-8489, Japan; Institute of Neuropsychiatry, 91 Bentencho, Shinjuku-ku, Tokyo 162-0851, Japan; Graduate School of Informatics, Kyoto University, 36-1 Yoshidahonmachi, Sakyo-ku, Kyoto 606-8501, Japan

## Abstract

The free-energy principle has been proposed as a unified theory of brain function, and recent evidence from in vitro experiments supports its validity. However, its empirical application to in vivo neuronal dynamics during active decision-making remains limited. This work reverse-engineered generative models—cast as canonical neural networks—from large-scale calcium imaging data of zebrafish performing a visually guided go/no-go task in a virtual-reality environment. Leveraging the formal equivalence between neural network dynamics and variational Bayesian inference, we constructed biologically plausible synthetic agents capable of active inference. These agents recapitulated individual variability in zebrafish neuronal dynamics and behaviour by identifying subject-specific prior and posterior beliefs. Additionally, they enabled quantitative predictions of long-term changes in neural activity, effective synaptic connectivity, and behavioural performance, including task accuracy after training. Our results demonstrate a powerful framework of active inference for modelling in vivo neuronal self-organisation and highlight the predictive validity of the free-energy principle in behaving animals.

## INTRODUCTION

Elucidating the computational principles underlying brain functions remains a central goal of neuroscience [1–4]. A fundamental criterion for any such principle is predictive power, that is, the ability to anticipate how brain function unfolds over time. Such predictions can deepen our understanding of the emergence of cognitive function and the progression of neurodegenerative and psychiatric disorders [5]. However, unlike recent successes in short-term predictions [6–9], predicting the long-term self-organisation of neural circuits—including synaptic plasticity and emergent network dynamics—remains challenging. This difficulty arises in part from the limited resolution and coverage of empirical measurements, which constrain bottom-up approaches that attempt to model neuronal self-organisation purely from data. To overcome these limitations, this work proposes the integration of empirically grounded inferences with normative computational theories of brain function.

The free-energy principle has emerged as a comprehensive theory of the brain to account for perception, learning, and action of all biological organisms [10,11]. The principle posits that the brain performs variational Bayesian inference [12] of external milieu by minimising variational free energy as a tractable proxy for minimising surprise. This entails the updates of posterior beliefs about external milieu states encoded by neural activity and synaptic weights. Moreover, animals can infer and then select actions that minimise expected risk associated with future outcomes, a process termed active inference [13,14]. These types of inferences rest upon generative models, that is, internal hypotheses or representations of the external milieu that express how external states generate sensory input. Recent theoretical work has shown that canonical neural networks can implement such generative models within their network architecture, thereby performing active inference through neural activity and synaptic plasticity in a biologically plausible manner [15–18]. In particular, delayed modulation of Hebbian plasticity [19–22] in the form of three-factor learning [23–25] is a possible implementation of active inference, in which agents minimise future risks by retrospectively updating behavioural strategies [16].

Despite numerous theoretical works, the free-energy principle has been criticised for its apparent unfalsifiability owing to the broad generality of its claims. However, it becomes testable when applied to specific systems with well-defined biological constraints [26]. To this end, previous works have developed a reverse engineering scheme to identify generative models from empirical neural data [15,16] and have demonstrated that this principle can accurately predict the self-organisation of in vitro neural networks of rat cortical cells when assimilating sensory information [27]. However, extending this approach to behaving animals in vivo remains a critical step toward validating the relevance of this principle to naturalistic cognitive function.

To pursue this type of validation, zebrafish are an ideal model animal owing to their genetic tractability and amenability to large-scale calcium imaging [28,29]. Despite their simplicity compared to mammals, zebrafish exhibit conserved neuroanatomical structures and behavioural capabilities [30–32]. Previous work established a go/no-go decision-making paradigm in a virtual reality environment, in which zebrafish engage in state inference and action selection by associating colour cues and visual flow with punishment, consistent with active inference [33]. Their rapid learning within a few hours facilitates continuous tracking of large-scale neuronal dynamics across learning phases, making them suited for comparing empirical changes with theoretical predictions [34].

In the present work, we reverse engineered canonical neural network models from zebrafish calcium imaging data during a decision-making task, yielding individual-specific generative models. These models enabled the construction of ‘synthetic fish agents’ that perform active inference, recapitulating the inter-individual variability in behavioural performance and neural circuit properties—including differences in internal representation, effective synaptic connectivity, and subjective risk between learners and non-learners. Subsequently, we demonstrated the predictability of these agents for unseen data by reverse engineering generative models based only on the initial empirical data before punishment was provided. These agents could predict long-term changes in neural activity and behaviour over the training period, outperforming conventional statistical and theory-based methods. Our results highlight active inference as a mechanistic and predictive framework for modelling in vivo neuronal self-organisation.

## RESULTS

### Zebrafish go/no-go decision-making paradigm

We began by outlining a zebrafish go/no-go decision-making task [33]. Head-fixed zebrafish were immersed in a virtual reality environment created using visual displays (**Fig. 1a**). Tail motion, captured by the camera, allowed the fish to navigate forward in the virtual space. Forward movement was visualised as black stripes moving against a coloured background. After an interval with a white background, the fish randomly engaged in either the go or no-go trial. In the go trial (**Fig. 1a**, top), the fish began in the blue zone; remaining there for 10 s triggered an aversive weak electric stimulus, whereas forward movement into the red zone prevented punishment. Conversely, in the no-go trial (**Fig. 1a**, bottom), fish started in the red zone; remaining there for 10 s prevented punishment, whereas moving forward into the blue zone was punished.

**Fig. 1.**
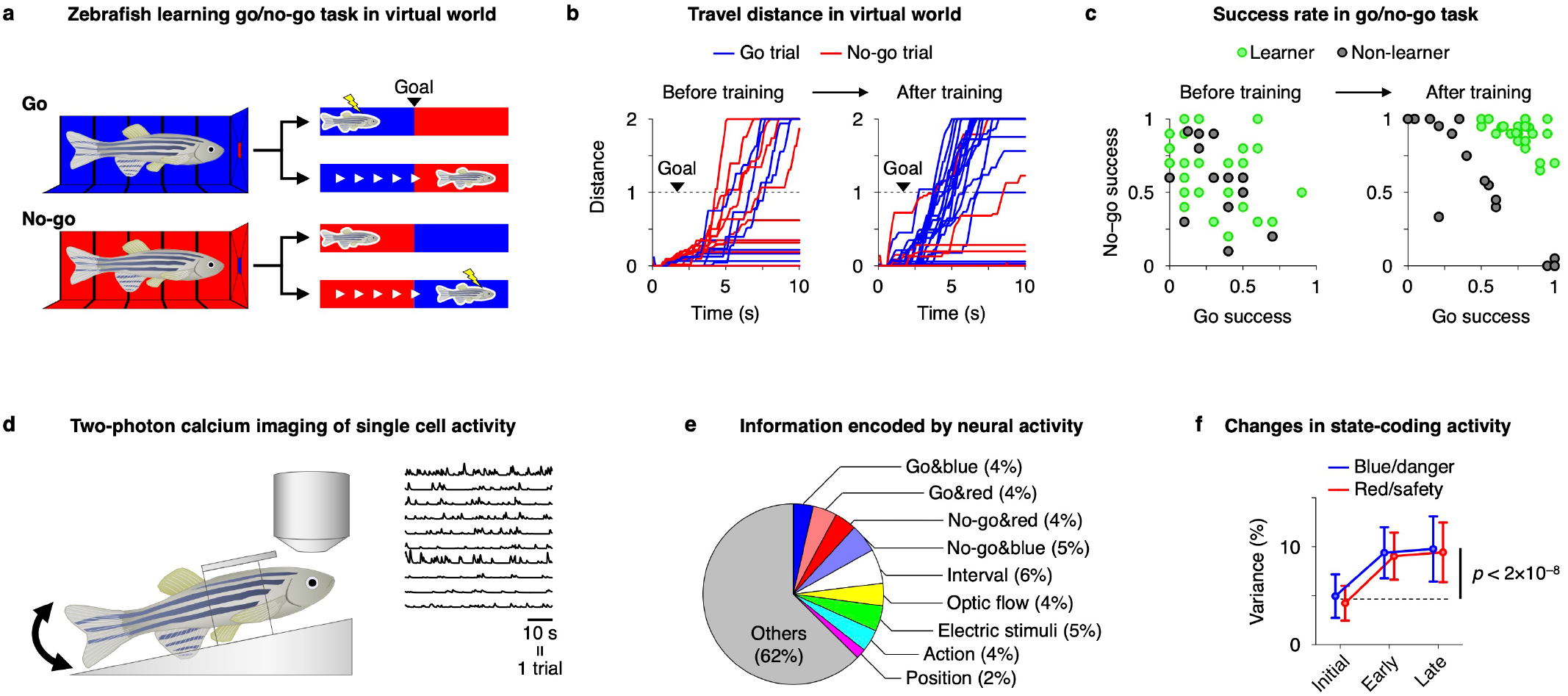
Zebrafish go/no-go decision-making paradigm. **a**. Schematic of a go/no-go decision-making task for zebrafish. Previous work developed a virtual reality-based experimental paradigm to train zebrafish [33], where head-fixed zebrafish was placed in a chamber surrounded by monitors to provide a virtual reality environment. Zebrafish underwent a go/no-go decision making to swim in response to blue cue (go, top) and remain in position in response to red cue (no-go, bottom) to avoid weak electric stimuli. **b**. Zebrafish movements before (left) and after (right) training. Data from a typical learner are shown. **c**. Task success rates before (left) and after (right) training. Data from 30 learners and 15 non-learners are shown. **d**. Two-photon calcium imaging of head-fixed zebrafish. Neural activity was measured during learning of the decision-making task across the telencephalon [33]. **e**. Pie chart representing the ratio of variances of neural activity explained by nine external information. The averaged contributions over 17,226 neurons from 39 fish are shown. **f**. Changes in neural activity encoding blue and red states. Their increase during training implies that these neurons self-organised to encode danger and safety states. Lines and error bars represent mean values ± SDs. In (**d–f**), a deconvolution filter was applied to the region of interest (ROI) signals, and 17,226 neurons from 39 fish that were properly detected activity were analysed. However, as the deconvolution filter was unable to properly extract activity from some neurons owing to noise, we used ROI signals without deconvolution in the subsequent analysis.

These randomly interleaved trials required the fish to infer the hidden environmental state— that is, whether the environment is safe or dangerous—based on sensory input and select their actions accordingly. During the initial 20-trial adaptation phase, no punishment was delivered, and the fish exhibited inherent (i.e., unconditioned) swimming behaviour irrespective of the conditions (**Fig. 1b**, left). Training continued for 80 trials and was divided into early (21–60) and late (61–100) phases. As the training progressed, the fish learned to swim selectively in the go trials and to remain still in the no-go trials (**Fig. 1b**, right). Learners were defined as individuals with success rates of at least 0.5 in both go and no-go trials and a marginal success rate of at least 0.6 during the late training phase (**Fig. 1c**). Of the 45 fish, 30 met these criteria and were classified as learners, whereas the remaining 15 were designated non-learners.

To investigate the neuronal basis of inference and decision-making in zebrafish, two-photon calcium imaging was performed during the aforementioned go/no-go task (**Fig. 1d**). From each fish, 440 ± 153 (mean ± standard deviation (SD)) neurons were recorded across the telencephalon, and they exhibited activity patterns indicative of internal representations of the external milieu. We first quantified the extent to which neural activity could be explained by task-relevant variables. Approximately 38% of the observed activity variance was accounted for by a set of nine sensory and behavioural features, including four conditions determined by two task types and colour cues (go&blue, go&red, no-go&red, and no-go&blue), inter-trial interval (white), optic flows, weak electric stimuli, motor actions, and position (**Fig. 1e**). Notably, neural activity encoding danger/safety states in response to colour cues (blue/red) occupied a substantial portion of the task-related neural variance, and the amplitudes of their variances increased during the training period (**Fig. 1f**). These results suggest that the zebrafish neurons formed internal representations of task-relevant hidden states and determined their behaviour accordingly.

### Reverse engineering generative models from neuronal data

The aforementioned observations were incorporated into the modelling as follows: in our task design, the input generative process was defined by two latent environmental states—danger and safety—represented as a two-dimensional hidden state vector *s*(*t*) = (*s*_*danger*_(*t*), *s*_*safety*_(*t*))^T^, and their transitions were determined by the fish’s action *δ*(*t*) (**Fig. 2a**, top). These hidden states generated four-dimensional sensory inputs *o*(*t*) = (*o*_*blue*_(*t*), *o*_*red*_(*t*), *o*_*flow*_(*t*), *o*_*stim*_(*t*)) ^T^, expressing blue, red, and flow visual inputs, and weak electric stimuli (**Fig. 2a**, bottom).

**Fig. 2.**
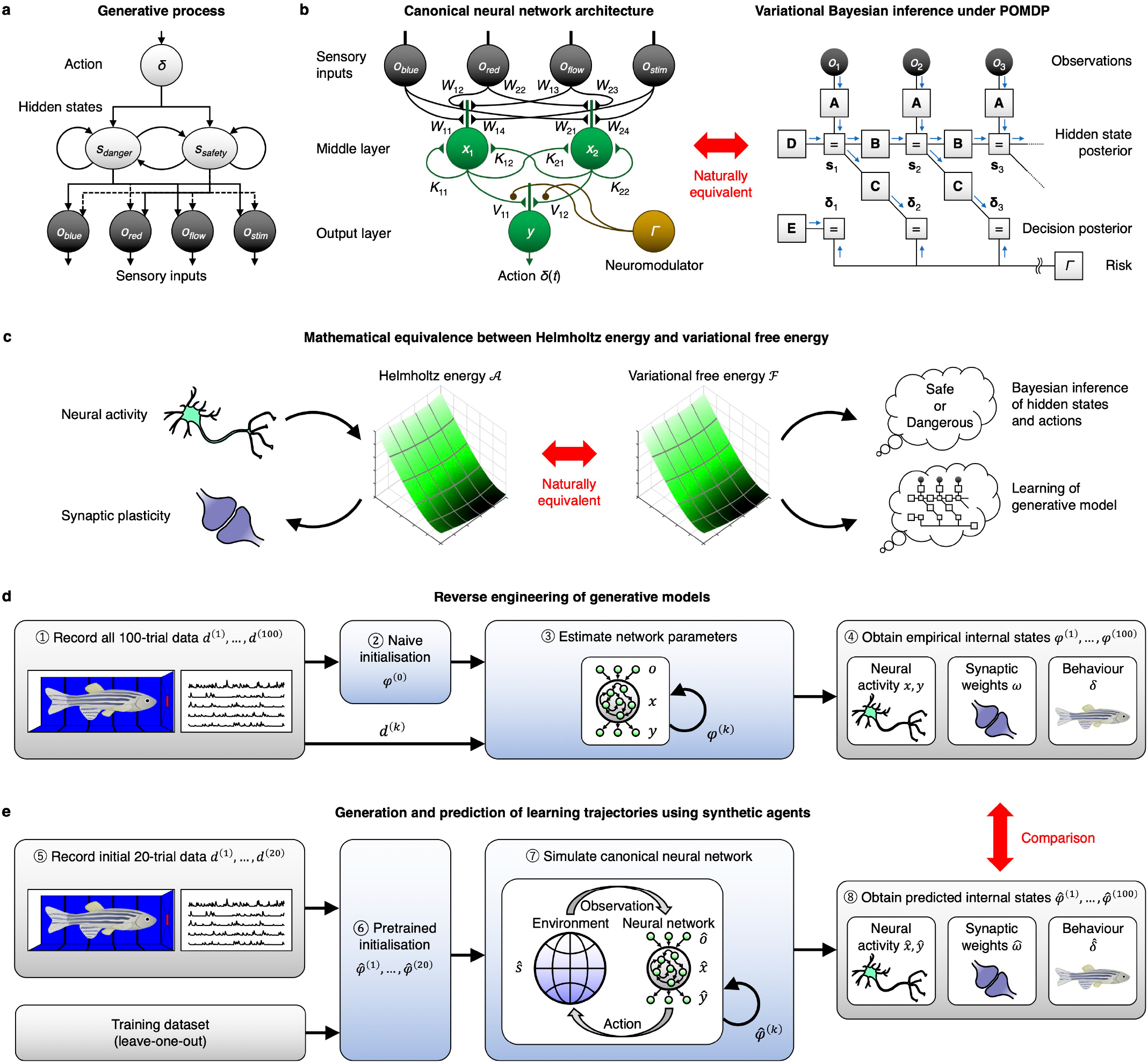
Validation of active inference using reverse engineering. **a**. Generative process for the considered go/no-go task. **b**. Canonical neural network of zebrafish. Left: Canonical neural network architecture. The middle and output-layer neural activities are expressed as *x*(*t*) and *y*(*t*). Right: The network dynamics can be read as performing Bayesian inference under a partially observable Markov decision process (POMDP) [52−54]. **c**. Mathematical equivalence between Helmholtz energy and variational free energy [15−18]. First, the neural activity equations for canonical neural networks are defined (top left). By computing the integral of these equations, biologically plausible Helmholtz energy 𝒜 can be reconstructed (middle left). The derivative of 𝒜 with respect to synaptic strengths yields equations of Hebbian plasticity (bottom left). Then, variational free energy ℱ is defined based on the generative model (middle right). Updating posterior beliefs about hidden states and parameters to minimise ℱ results in Bayesian inference (top right) and learning (bottom right). As shown in the middle, 𝒜 and ℱ have the identical functional form, as depicted by one-to-one correspondences between their components [16] (see also **Table 1** and the Methods section). **d**. Procedures for reverse engineering generative models using empirical neural activity data. Provided with empirical data *d*^(*k*)^ for trials *k* = 1, …, 100, unknown network parameters—including matrices for categorising neuronal ensembles, effective synaptic weights, firing thresholds, and subjective risks—are estimated by minimising 𝒜. **e**. Procedures for applying the reverse engineering method to predict learning trajectories. Unlike (**d**), here, reverse engineering was conducted using only the initial empirical data to construct synthetic agents employing individual-specific generative models. These agents can derive plausible synaptic plasticity rules and perform active inference to make the quantitative prediction of subsequent learning, enabling forward inference or extrapolation of neural activity, synaptic weights, and actions throughout training. Further details are provided in the Methods section.

Then, we modelled zebrafish neuronal networks using canonical neural networks that offer biologically plausible dynamics consistent with established realistic neuron models [16] (**Fig. 2b**, left). The network comprises two middle-layer neuronal ensembles *x*(*t*) = (*x*_1_ (*t*), *x*_2_ (*t*))^T^, which receive sensory inputs *o*(*t*) and produce an output-layer activity *y*(*t*). The output activity *y*(*t*) drives the agent*’*s actions *δ*(*t*) (i.e., tail motions). These neural activities follow differential equations for rate-coding models with sigmoidal activation functions derived as a gradient descent on Helmholtz energy 𝒜 (**Fig. 2c**, top left; refer to the Methods section for equations). The changes in synaptic weights (denoted as matrices *W, K*, and *V*) follow conjugate synaptic plasticity rules derived as a gradient descent on the same 𝒜 with respect to these weights, thus furnishing biologically grounded learning rules (**Fig. 2c**, bottom left). In particular, the update of the output-layer synaptic weights *V* = (*V*_11_, *V*_12_) takes the form of a three-factor learning rule [23−25], which is mediated by risk-encoding neuromodulators *Γ* such as dopaminergic or noradrenergic neurons [16].

**Table 1.**
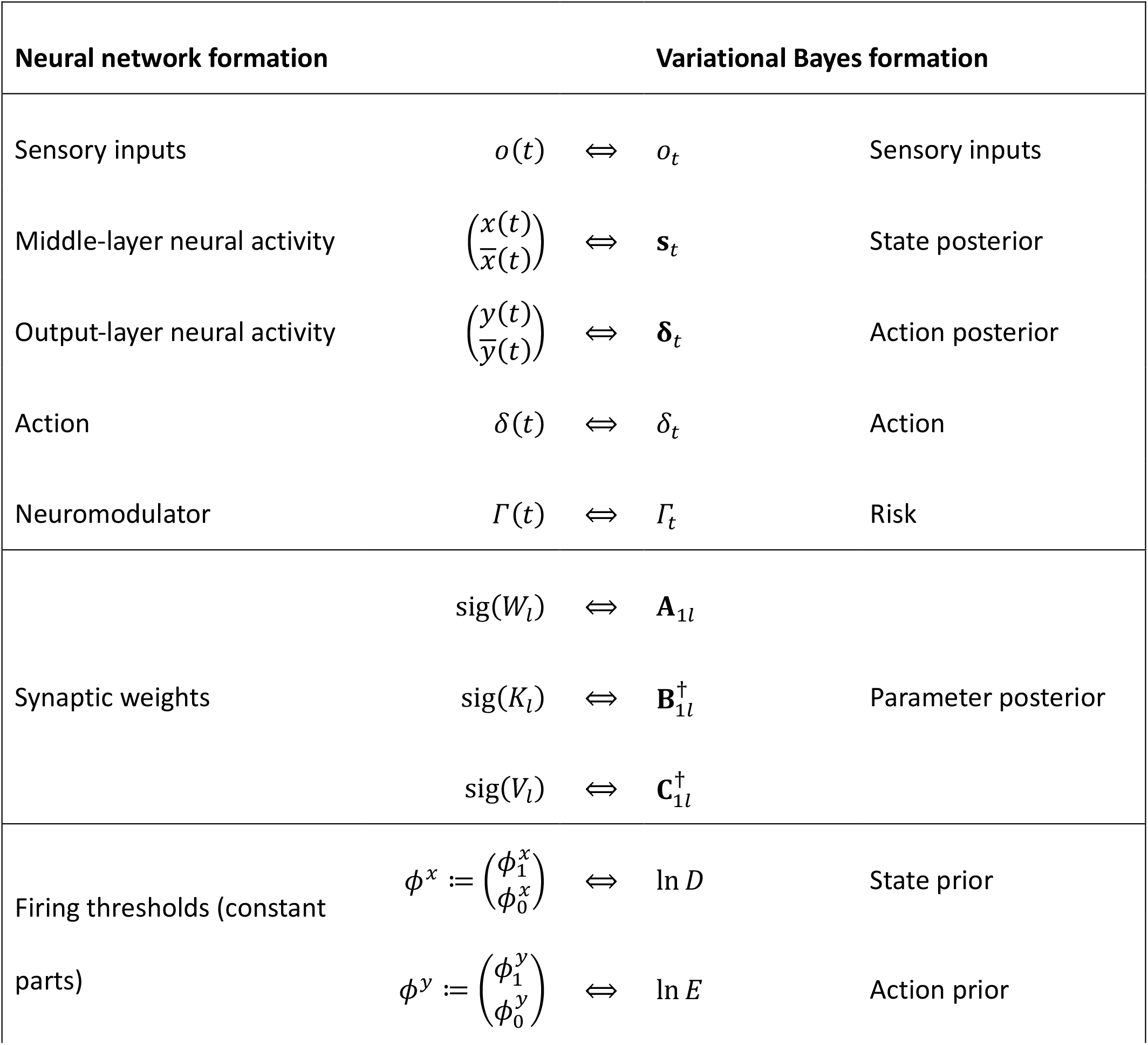
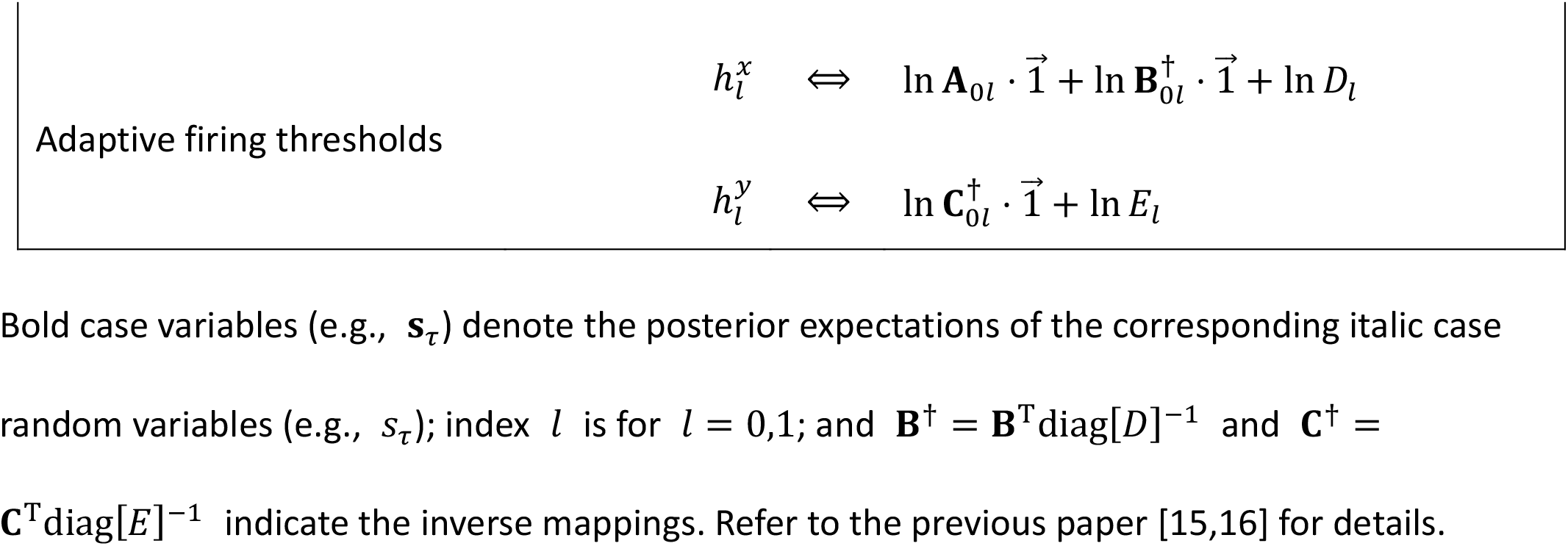
Correspondence of variables and functions.

Previous works have mathematically illustrated a formal equivalence between the Helmholtz energy 𝒜 in canonical neural networks (**Fig. 2c**, middle left) and variational free energy ℱ in the free-energy principle (**Fig. 2c**, middle right) by showing one-to-one correspondences between their components [15–18] (**Table 1**), that is, 𝒜 ≡ ℱ. This mathematical equivalence indicates that the neural activity and synaptic plasticity that minimise the shared Helmholtz energy can be conceptualised as performing active inference, that is, Bayesian belief updating to infer latent environmental states and actions (**Fig. 2b**, right). For instance, delayed modulation of Hebbian plasticity [19–22] is shown to be a sufficient neuronal substrate for implementing active inference that minimises risk associated with future outcomes [16]. This equivalence offers a unified explanation for neuronal dynamics and computation, providing a normative and mechanistic account that links physiological phenomena to cognitive functions.

Building on this foundation, we propose a method to reverse-engineer generative models from empirical brain activity data and test the predictive validity of active inference [26,27]. The procedure begins by fitting experimentally recorded neural activity to a canonical neural network (**Fig. 2d**, left), in which empirical neural activities were systematically allocated to the two ensembles *x*(*t*) by minimising 𝒜. This allocation is a projection of high-dimensional neuron-wise data onto a two-dimensional subspace of *x*(*t*) that maximises predictability, thus facilitating the extraction of task-relevant neural activity. From this, latent internal parameters—including effective synaptic connectivity, firing thresholds, and subjective risks—were statistically estimated by minimising the same 𝒜 (**Fig. 2d**, right). A complete description of the procedure is provided in the Methods section “Reverse engineering of generative models”.

Once the canonical neural network has been reconstructed, the associated Helmholtz energy 𝒜 can be used to derive the corresponding variational free energy ℱ and its generative model (**Fig. 2c**). This enables a formal, one-to-one mapping between neural network properties and those of Bayesian inference under a certain generative model that embodies an animal’s internal hypotheses (**Table 1**). Under this framework, empirical properties of neural activity and synaptic weights were made explainable in terms of posterior beliefs, enabling a formal characterisation of the animal’s perceptions and actions.

Subsequently, empirically derived generative models are used to construct synthetic agents capable of prediction, learning, and action selection (**Fig. 2e**). The gradient descent on ℱ furnishes a synaptic plasticity rule for these synthetic agents. Crucially, this framework supports forward inference (i.e., extrapolation): the free-energy principle posits that neural activity and synaptic plasticity pursue the free energy gradient. Therefore, if active inference is applied, these synthetic agents should be able to predict future latent variables and emulate an animal’s behaviour and neuronal dynamics when exposed to the same task environment. This allows us to test the predictive validity of the free-energy principle and active inference [26].

In the remainder of this paper, we applied this analytical paradigm to reverse engineer the generative models underlying zebrafish decision-making. We then examined whether active inference instantiated in a canonical neural network could quantitatively predict the behaviour, brain activity, and learning dynamics of individual animals.

### Plausibility and individual variability in zebrafish generative models

Here, reverse-engineered generative models were analysed to verify some qualitative predictions of the free-energy principle [10,11]. Ensemble *x*_1_ received dominant sensory input from the blue and flow cues and was highly active during the go trials, consistent with encoding a danger-related hidden state (**Fig. 3a**, left; data from *n* = 30 learners). In contrast, ensemble *x*_2_ was excited by the red input and was more active in the no-go trials, suggesting its role in encoding the safety state (**Fig. 3a**, centre). Upon receiving the inputs from these ensembles, larger motor outputs were produced in the go trials (**Fig. 3a**, right). These functional specialisations emerged during task acquisition, as evinced by the progressive increase in correlations between ensemble activities and hidden states (danger/safety) during training (**Fig. 3b**). Crucially, ensemble *x*_1_ integrated multimodal sensory information, not only colour but also optic flow (**Fig. 3c**), highlighting their role in inferring hidden states through evidence accumulation. These dynamics are consistent with in vitro empirical support of the free-energy principle [27,35,36] and in vivo evidence of Bayesian inference [37,38] that show that neural activity self-organises to encode hidden states or latent causes of sensory inputs.

**Fig. 3.**
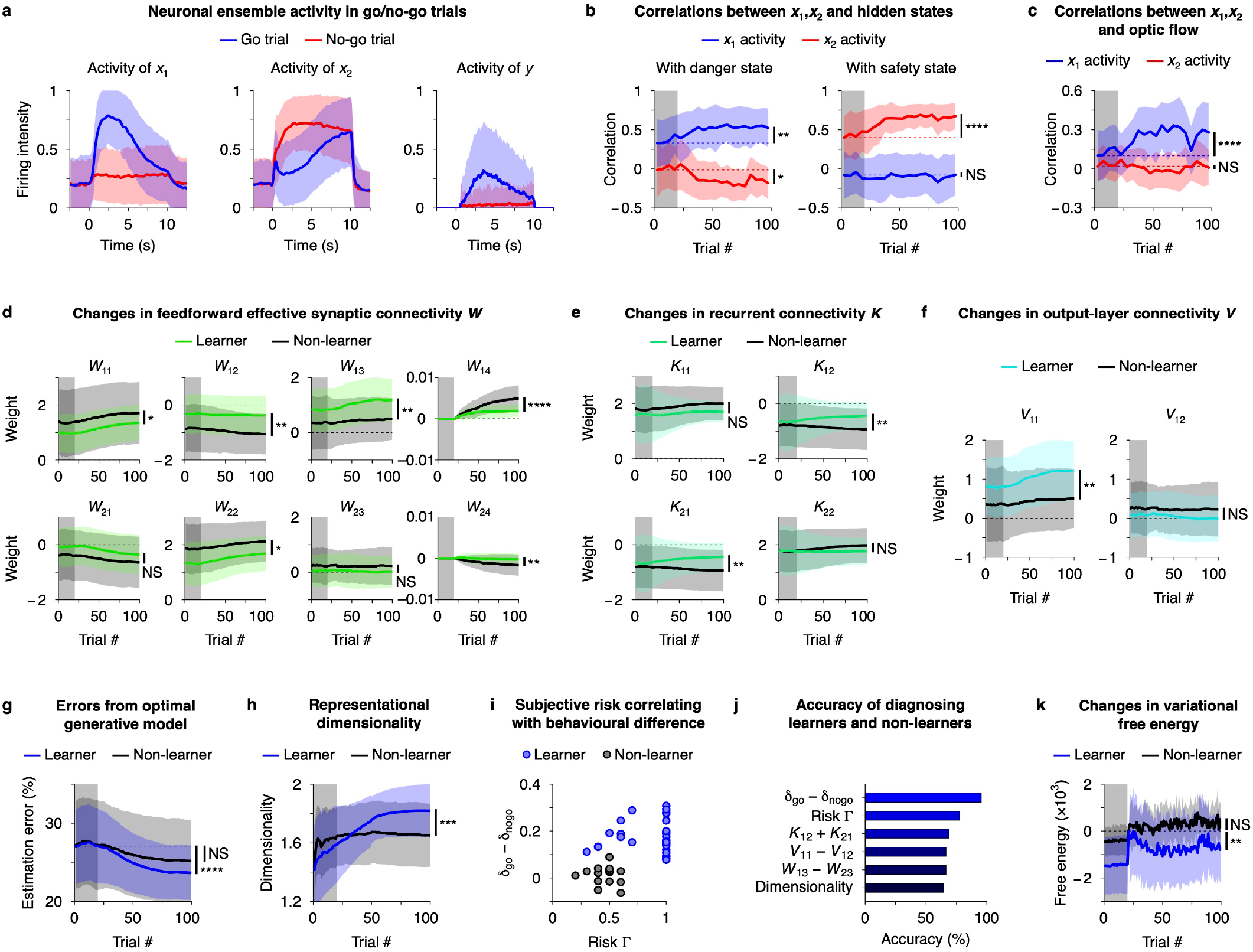
Reverse-engineered zebrafish generative models. **a**. Neuronal ensemble activities in the go (blue) and no-go (red) trials during the late training period. In (**a–c**), data obtained from *n* = 30 learners were used. **b**. Changes in correlations between ensemble activity *x*_1_, *x*_2_ and hidden danger/safety states over trials. **c**. Changes in correlations between *x*_1_, *x*_2_ and optic flow input. **d− f**. Changes in the effective synaptic connectivities in the middle (*W, K*) and output (*V*) layers. Comparisons between trajectories of learners (coloured, *n* = 30) and non-learners (black, *n* = 15) are shown. Here, *W*_*i*1_, *W*_*i*2_, *W*_*i*3_, and *W*_*i*4_ denote forward effective connections from blue, red, flow visual cues and weak electric stimuli to ensemble *x*_*i*_, respectively; *K*_*ij*_ denotes a recurrent connection from ensemble *x*_*i*_ to *x*_*j*_; and *V*_1*i*_ denotes a connection to the output (*i, j* = 1,2). **g**. Reduced difference between the empirical network architecture and the optimal generative model. The mean squared errors between parameter posteriors in the empirical generative model and theoretical values of *A, B*, and *C* matrices are plotted (see the Methods section for definitions of *A, B*, and *C*). **h**. Significant differences in the representational dimensionality of neuronal ensemble activity between learners and non-learners. High representational dimensionality indicates functional segregation of the two ensembles. **i**. Correlation between estimated subjective risk (*Γ*) and behavioural difference between go and no-go trials (i.e., *δ*_*go*_ *− δ*_*nogo*_) during the late training period. The subjective risks for receiving punishment were estimated using Bayesian model selection. **j**. Diagnosis of learners and non-learners using network properties. **k**. Variational free energy computed from data decreasing with trials. Changes from the first training trial (trial 21) are shown. In (**d−k**), data from *n* = 30 learners and *n* = 15 non-learners are compared. Lines and shaded areas in (**a−h, k**) represent mean values ± SDs. NS, *p* ≥ 0.05; *, *p* < 0.05; **, *p* < 0.01; ***, *p* < 0.001; ****, *p* < 0.0001. Grey areas in (**b–h, k**) indicate the initial 20-trial adaptation period, in which no punishment was delivered. Further details are provided in the Methods section.

These changes result from activity-dependent synaptic plasticity. Effective synaptic connectivity analysis revealed distinct network properties between learners and non-learners (**Fig. 3d–f**). In particular, the effective synaptic weight from the flow input to *x*_1_ (*W*_13_) increased selectively in learners during training (**Fig. 3d**), suggesting that flow information is critical for decision-making in successful fish. The network architecture in learners was also characterised by minimal recurrent inter-ensemble connections (*K*_12_, *K*_21_), which promoted the functional segregation of the two ensembles (**Fig. 3e**). By contrast, non-learners formed stronger inhibitory inter-ensemble connections during training, which reflected a disrupted representation of the task structure.

Learner networks further exhibited strengthened positive connections from *x*_1_ to the output (*V*_11_), which facilitated action initiation in the go trials, whereas minimal connections from *x*_2_ (*V*_12_) suppressed actions in the no-go trials (**Fig. 3f**). This asymmetry created a positive feedback loop linking blue-and-flow perception to action, enabling continuous movement during the go trials.

The learner network architecture became increasingly closer to that of the optimal generative model to minimise future risks in the given task setting, as predicted by the free-energy principle (**Fig. 3g**). Learners also developed more differentiated and richer internal representations of hidden environmental states, which were reflected in the higher effective dimensionality [39] of the ensemble activity that was close to two dimensions (**Fig. 3h**). The learning inability in non-learners can be attributed to reduced representational dimensionality (**Fig. 3h**) and considerable inhibitory coupling (negative correlation) between neural ensembles (**Fig. 3e**), which might be caused by strong initial inhibitory inputs from minor colour cues such as *W*_12_ and *W*_21_ (**Fig. 3d**). These features are associated with strong prior beliefs, which are consistent with previous theoretical [15] and in vitro findings [27,35]. These results suggest that learners placed importance on sensory evidence, whereas non-learners placed more importance on their prior beliefs.

Moreover, we assessed individual differences in punishment-related learning by estimating the subjective risk *Γ* for each fish (**Fig. 3i**). Learners exhibited higher perceived risk (mostly close to one) in response to punishment, which enhanced the learning of output-layer weights (*V*) to avoid adverse outcomes, enabling sharper behavioural discrimination. Conversely, non-learners exhibited low subjective risk (approximately 0.5) and weak behavioural contrast between trial types, implying insufficient internal modelling of the task structure. These differences highlight the intrinsic difference in their behavioural optimisation criteria.

These correlations between the network properties and task success can be used for the diagnosis of learners and non-learners (**Fig. 3j**). The differences in mean actions within the go and no-go trials, subjective risks, and synaptic weights were highly predictive of task success. A subset of fish showed high representational entropy but poor task performance, suggesting a two-stage failure mode, that is, inadequate internal modelling or impaired action selection. Both can be attributed to differences in prior beliefs that deviate the generative model from the Bayes-optimal form.

Furthermore, we confirmed the reduction of variational free energy ℱ over training for learners (**Fig. 3k**). Learners exhibited a larger reduction in ℱ than non-learners, indicating better alignment of their generative models with the task environment. These results directly support the notion that learning in zebrafish conforms to the free-energy principle.

In essence, individual differences in zebrafish learning were attributed to variations in the generative model structure implicit in canonical neural networks. The ability to infer and act upon hidden states relies on how sensory evidence and prior beliefs are integrated, which determines the self-organisation of internal representations over time. These findings provide a mechanistic foundation for understanding cognitive variability and predictive processing in vertebrate brains and a basis for developing long-term prediction models.

### Quantitative prediction of zebrafish learning using synthetic agents

Next, we examined the predictive validity of active inference by asking whether synthetic agents—characterised by initial neural data from individual zebrafish—could predict both learning-related changes in neuronal dynamics and behavioural performance (**Fig. 4a**). Despite the inherent difficulty of long-term prediction in biological systems, our analyses revealed that key differences between learners and non-learners could be traced back to the initial network parameters (**Fig. 3d–f**). In particular, individuals with strong red-inhibitory (*W*_12_) and weak flow-excitatory (*W*_13_) initial weights resulted in a failure to acquire task-relevant representations, identifying these individuals as non-learners (**Fig. 3d**). These observations imply that learning trajectories can be predicted from neural activity measured during the initial adaptation period.

**Fig. 4.**
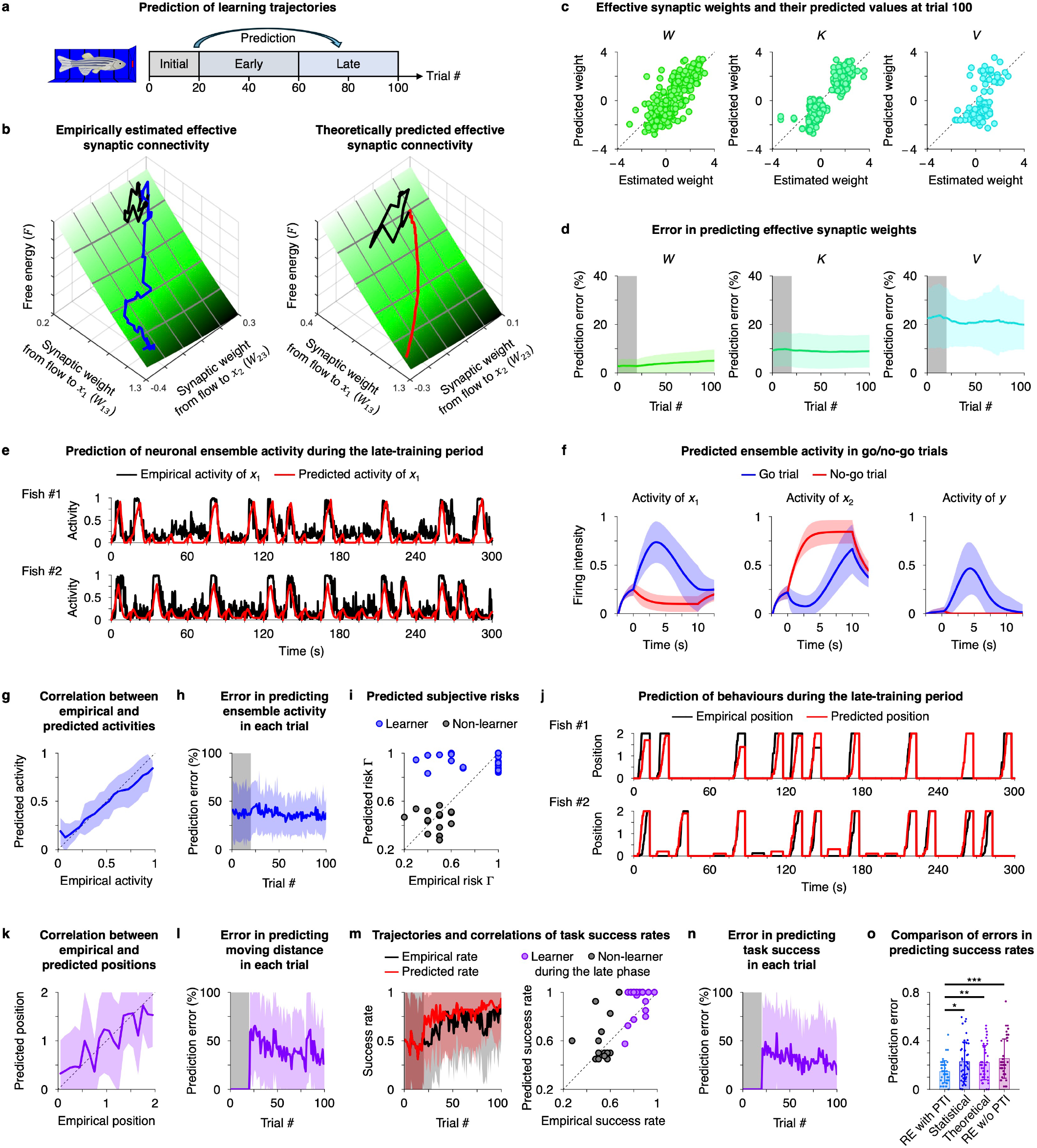
Quantitative predictions of zebrafish self-organisation. **a**. Schematic for long-term prediction. The synthetic agents reverse engineered based on empirical data in the initial 20-trial adaptation period are used to predict individual learning trajectories in biological fish during early and late training phases up to trial 100. **b**. Comparison of empirical results and theoretical predictions. Left: Effective synaptic weights (*W*_13_, *W*_23_) estimated from the data of a representative learner. Black and blue lines are trajectories during trials 1–20 and 21–100, respectively. Right: Predicted weights in the absence of data (red). **c**. Comparison of the estimated and predicted effective synaptic weights at trial 100. **d**. Errors in predicting effective synaptic weights. The errors are assessed based on the mean squared error between estimated and predicted weights, divided by the squared amplitude of weights. (**c, d**) are obtained from *n* = 360, 180, and 90 connections for *W, K*, and *V*, respectively. **e**. Empirical and predicted activities of ensemble *x*_1_. Concatenated activity trajectories during trials 61−80 after excluding parts of rest period data are shown. **f**. Predicted ensemble activities obtained in the go (blue) and no-go (red) trials during the late training phase. **g**. Correlations between empirical and predicted activities during the late phase. **h**. Error in predicting ensemble activity. The ratio of squared prediction error and squared amplitude is plotted. **i**. Predicted subjective risks estimated using the pretrained initialisation function. **j**. Empirical and predicted moving distance in trials 61−80. **k**. Correlations between empirical and predicted moving distances (positions). **l**. Error in predicting moving distance. **m**. Trajectories of empirical and predicted success rates (left) and their correlations during the late phase (right). **n**. Error in predicting task success in each trial, measured based on the absolute error. **o**. Comparison of task-success-rate prediction errors between reverse engineering with pretrained initialisation (RE with PTI) and three conventional methods: naive statistical method using single-layer perceptrons, theory-based method using canonical neural networks, and reverse engineering without pretrained initialisation (RE w/o PTI). In (**c, d, i, m−o**), data from *n* = 30 learners and *n* = 15 non-learners were used. In (**f−h, k, l**), data from *n* = 30 learners were used. (**b, e, j**) are data from representative learners. Lines and shaded areas in (**d, f–h, k–n**) represent mean values ± SDs. Grey areas in (**d, h, l–n**) indicate the initial 20 trials. *, *p* < 0.05; **, *p* < 0.01; ***, *p* < 0.001. Further details are provided in the Methods section.

To enable data-driven reverse engineering that incorporates individual fish properties, we developed a pretrained initialisation function designed to estimate the initial network parameters (**Fig. 2e**; refer to the Methods section “Prediction of learning trajectories” for details). Trained on an activity dataset of the other fish, this function estimates the initial internal variables for each fish—including the initial synaptic weights and subjective risks—based on the neural activity data during the initial 20-trial adaptation period (**Fig. 2e**, left). Throughout all predictive analyses, we employed a leave-one-out cross-validation approach [40] at the individual fish level: for each test fish, hyper parameters of the initialisation function were optimised using the entire dataset from all other fish. Then, the test fish’s learning trajectory was predicted based only on initial 20-trial data of the fish using the pretrained function and canonical neural network simulations (**Fig. 2e**, right), ensuring that no information from future trials was used during predictions.

Consistent with the free-energy principle [10,11], the plasticity in the estimated synaptic weights occurred in the direction of minimising the variational free energy ℱ along its gradient, as illustrated in the (*W*_13_, *W*_23_) trajectories (**Fig. 4b**, left). This provided direct evidence that synaptic plasticity in zebrafish neural circuits conforms to the normative Bayesian belief updating rules. Based on these observations, we constructed synthetic zebrafish agents whose canonical neural networks performed active inference. They employed generative models estimated from neural activity data during the initial 20-trial adaptation period. We then exposed these agents to the same go/no-go task environment as biological fish for training. These synthetic agents— identical to ideal Bayesian observers [15,16]—learned by pursuing a gradient descent on ℱ, forming posterior beliefs about latent danger/safety states and selecting actions accordingly. We confirmed that the predicted synaptic weight trajectories matched those estimated from empirical data (**Fig. 4b**, right), demonstrating that the free-energy gradient was a key aspect to characterise the plasticity direction.

We then quantitatively assessed the accuracy of these predictions. The predicted synaptic connectivity changes in both the middle and output layers aligned well with the experimentally estimated values (**Fig. 4c**). Across fish, the synaptic weights predicted after training deviated by approximately 5%, 10%, and 20% from the empirical measurements for *W, K*, and *V*, respectively, confirming the precision of the model in capturing learning trajectories (**Fig. 4d**). In particular, the risk-modulated synaptic plasticity in the output layer (*V*) could recapitulate empirical observations of task-dependent plasticity in zebrafish. Therefore, the synthetic active inference agents successfully recapitulated zebrafish learning.

Moreover, the synthetic agents could predict neural activity patterns over training (**Fig. 4e–h**). Importantly, this constitutes a highly nontrivial prediction, as synaptic efficacy is dynamically modulated by activity, and neuronal responses are themselves contingent upon synaptic weights. Although activity predictions were more variable than synaptic weights—owing to intrinsic noise in the neural data—the overall structure of the inferred representations was preserved (**Fig. 4e**). The predicted activity patterns (**Fig. 4f**) were homologous to empirical activity (**Fig. 3a**), and these activities were correlated at each trial (**Fig. 4g**; Pearson correlation coefficient *r* = 0.76), even during the late training period. The error in predicting ensemble activities remained approximately 35% during the late training period (**Fig. 4h**), indicating that despite noise and variability, the model successfully captured core features of the learning process.

Additionally, behavioural metrics such as moving distance and success rate were predicted based solely on the initial neural activity. Based on the initial synaptic weights, we derived the predicted values of subjective risk *Γ* before fish has received punishment (refer to the Methods section for details) and confirmed the matching between empirical and predicted risks (**Fig. 4i**; *r* = 0.66), enabling us to simulate individual-specific punishment-related learning. The predicted behaviours (i.e., moving distances) in go and no-go trials were homologous to empirical ones (**Fig. 4j**) and correlated at each trial (**Fig. 4k**; *r* = 0.63). Interestingly, the errors in predicting positions tended to decrease as fish learned the task as behaviours of learned fish were more consistent with Bayes-optimal synthetic agents (**Fig. 4l**). Consequently, synthetic agents successfully predicted task success rates of biological fish over the training period (**Fig. 4m**; *r* = 0.77 during the late phase), and the prediction errors exhibited a decreasing trend during training (**Fig. 4n**). These results demonstrated that individual learning outcomes—including success rate and distance travelled—could be predicted from initial neural states using active inference.

We further benchmarked our reverse engineering framework with pretrained initialisation against existing approaches (**Fig. 4o**). Conventional statistical methods driven by empirical data lack mechanistic grounding, whereas theory-based Bayesian models—while offering qualitative insights—typically assume idealised parameters that limit individual-level predictions, both of which exhibited poor long-term predictions of task success rates (**Fig. 4o**, the centre two). The performance of reverse engineering without pretrained initialisation was also poor owing to the failure of capturing individual initial states (**Fig. 4o**, right). By contrast, the present method with pretrained initialisation explicitly incorporated biological variability by estimating individual-specific generative models, enabling more accurate long-term forecasts (**Fig. 4o**, left).

Taken together, our findings demonstrate that reverse-engineered synthetic agents can quantitatively predict individual zebrafish learning trajectories, highlighting the predictive validity of active inference in a biologically realistic setting. These agents quantitatively predicted key features of learning, including synaptic plasticity, neural activity pattern, and behavioural outcomes, based solely on initial empirical data. These results support the biological plausibility of the free-energy principle as a normative theory of brain function, providing a mechanistically explainable foundation for long-term predictions of brain self-organisation.

### Generalising predictions to novel task environments

So far, we demonstrated that our synthetic agents—reverse engineered from zebrafish neural activity—can quantitatively predict long-term learning trajectories in task settings identical to training settings. However, it remains unclear whether these agents can generalise their predictive capacities to unfamiliar environments that differ from the original training context. To address this, we evaluated model generalisation under two untrained conditions: an open-loop setting and a reversal learning paradigm (**Fig. 5**). Synthetic agents that had been trained under standard task contingencies were exposed to these conditions and compared with empirical data from zebrafish.

**Fig. 5.**
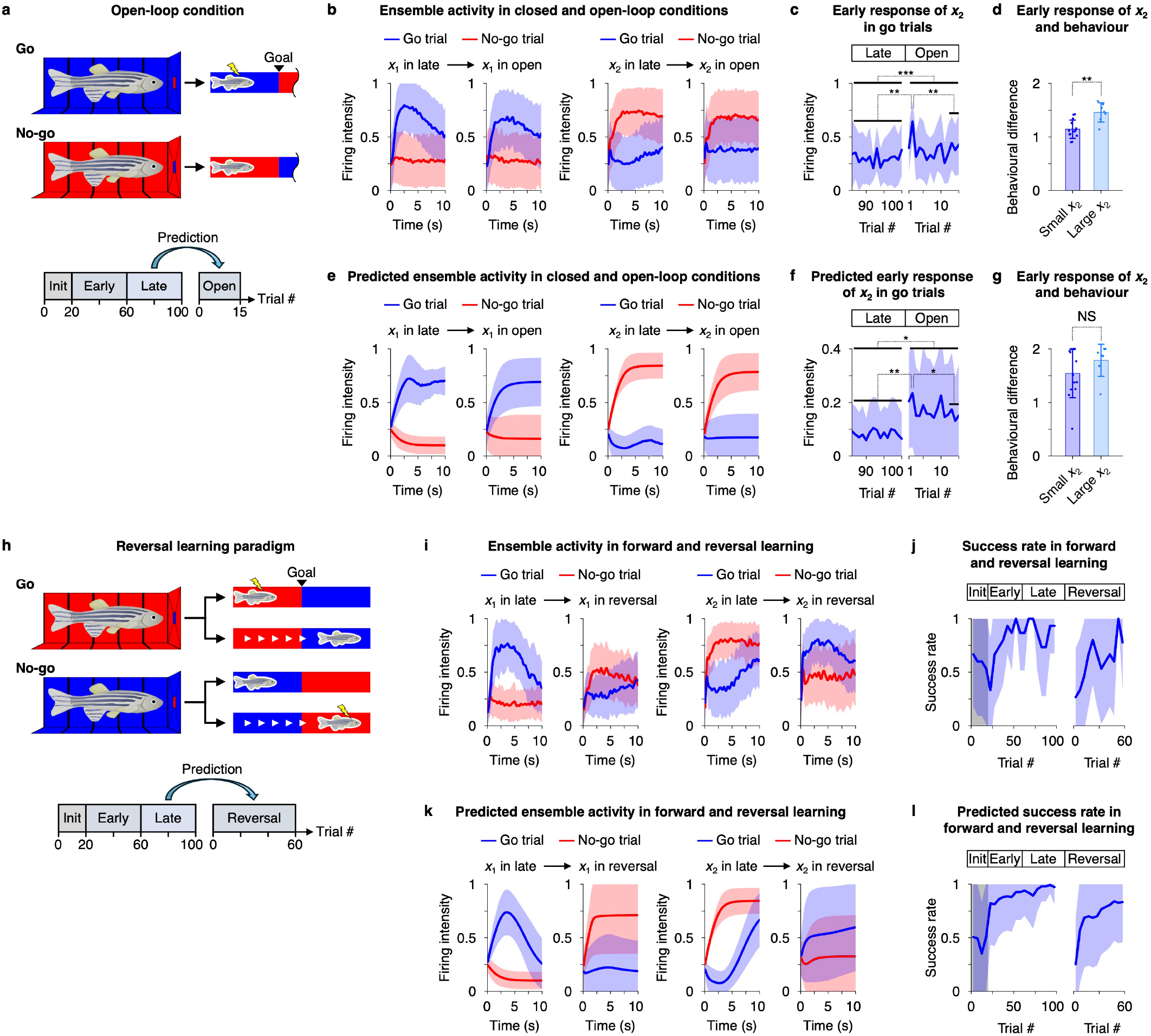
Generalising predictions to different task conditions. **a**. Schematic of open-loop conditions, in which neural outputs do not induce forward movement. **b**. Empirically observed activity during the closed-loop (trials 61–100 in the late phase) and open-loop (additional trials 1–15) conditions. The mean activities when observing blue (or red) cues in the go (or no-go) trials are plotted. In (**b– d**), data from *n* = 20 learners exposed to the open-loop condition are used. **c**. Changes in early response of *x*_2_ in the go trials when observing blue cues. **d**. Comparison of behavioural differences between learners with large *x*_2_ in the open-loop condition (*n* = 7, top 33%) and those with small *x*_2_ (*n* = 13). Behavioural differences are defined as the gap between mean actions in go and no-go trials during the late phase. **e**. Predicted activities of synthetic agents. Agents derived from *n* = 30 learners are used for simulations. The network’s plasticity insensitivity was reduced by half following initial task learning to facilitate plasticity. **f**. Predicted changes in early response of *x*_2_. **g**. Comparison of predicted behavioural differences between learners with large and small *x*_2_ during the open-loop condition. Results from 20 agents corresponding to the fish in (**g**) are shown. **h**. Schematic of reversal learning paradigm, in which the contingency between the color cues and punishment is inverted. **i**. Empirically observed activity during forward (trials 61–100 in the late phase) and reversal (additional trials 1–60) learning phases. The mean activities in go and no-go trials are plotted. In (**i, j**), data from *n* = 3 learners exposed to the reversal learning condition are used. **j**. Changes in task success rate during forward and reversal learning conditions. **k**. Predicted activities of synthetic agents. Agents derived from *n* = 30 learners are used for simulations. The network’s plasticity insensitivity was reduced following initial task learning and the three-factor learning rule mediated by risk *Γ* was applied to *W* during the reversal condition to facilitate plasticity. **l**. Predicted changes in task success rate. Lines and shaded areas (or error bars) in (**b−g, i−l**) represent mean values ± SDs. Grey areas in (**j, l**) indicate the initial 20 trials. NS, *p* ≥ 0.05; *, *p* < 0.05; **, *p* < 0.01; ***, *p* < 0.001.

In the open-loop condition, the fish tail movements no longer induced forward movement within the virtual environment (**Fig. 5a**). In this scenario, fish that had previously learned to escape from the danger state continued to attempt escape via tail beating, without success. Consistent with our previous work that reported elevated optic-flow prediction errors in mismatched sensorimotor contingencies [33], ensemble *x*_2_ displayed a persistent increased activity in the open-loop go trials, compared to the activity in the closed-loop go trials during the late training phase (**Fig. 5b**). These persistent activities associated with prediction errors were highest when the fish encountered the open-loop condition for the first time, then gradually decreased over the subsequent 15 trials—a pattern consistent with the free-energy principle (**Fig. 5c**). We also confirmed that learners with large increases in *x*_2_ during the open-loop conditions exhibited more efficient behaviours during the closed-loop conditions (**Fig. 5d**), consistent with our previous findings [33]. In addition to confirming these neural signatures, the synthetic agents could reproduce the deviated activities in the open-loop conditions (**Fig. 5e**) and forecast a gradual decline in these activities over time (**Fig. 5f**) and circuit-behaviour relationships (**Fig. 5g**), even in unfamiliar scenarios.

Next, we examined reversal learning, wherein the previously learned stimulus–response contingencies were inverted; that is, fish that had learned to swim when presented with a blue background now had to learn to swim in response to red (**Fig. 5h**). Initially, the fish failed to adapt, consistent with their prior beliefs. However, with repeated exposure, they gradually learned the reversed contingency by changing neural activity (**Fig. 5i**), as evinced by a gradual re-increase of task success rate (**Fig. 5j**). These reversed neural activity patterns in the reversal learning corroborated that ensembles *x*_1_ and *x*_2_ encoded the danger and safety states. The synthetic agents could reproduce these changes in neural activity (**Fig. 5k**). As expected, the behavioural trajectories predicted by the model closely matched the empirical reversal learning dynamics of biological fish (**Fig. 5l**). These results demonstrated that the reverse-engineered generative model—trained in a specific task environment—retained the capacity to generalise its predictions to novel conditions that were not used during training.

Therefore, our findings show that the inferred generative models are capable of recapitulating zebrafish learning and behaviour even under unfamiliar task variations. The resulting synthetic agents can effectively generalise predictions to new, untrained conditions, indicating that they capture core learning mechanisms of zebrafish. This generalisation capacity supports the robustness of the free-energy principle and active inference as an explanatory framework for adaptive behaviour in biological systems.

## DISCUSSION

Predicting whether an individual will successfully learn a task—or more broadly, whether they will develop into high-performing individuals—is of central interest in the practical applications of neuroscience. This challenge lies at the heart of our efforts to understand the neuronal mechanisms of learning and behaviour, with implications for psychiatry, education, and neuromorphic engineering. While such predictions are often intractable in complex systems, this work demonstrates that under well-defined conditions, it is possible to predict whether a zebrafish will become an effective learner or remain a non-learner solely based on its initial neuronal states.

The key innovation for this achievement is the incorporation of a normative theory of brain function—the free-energy principle [10,11]—into data-driven predictions. By applying the free-energy principle to in vivo neural recordings, we confirmed the predictive validity of this theoretical framework in behaving animals. We specifically demonstrated that active inference— when grounded in empirical data—can generate long-term predictions of individual learning trajectories. While previous works have largely focused on short-term predictions of brain activity, such as subsecond-scale electrophysiological signals [6–9], our approach enables forecasting of learning outcomes and neural self-organisation over training periods. This is also complementary to previous work that have predicted behavioural outcomes using machine learning approaches [41].

A central contribution of this work is the reverse engineering of generative models from empirical neural activity. In contrast to conventional statistical and machine learning approaches that primarily estimate static network architectures [42,43], our method reconstructs the underlying generative models that shape both neural dynamics and plasticity. Moreover, this framework enables data-driven identification of individual generative models, instead of explicitly assuming specific Bayesian models a priori, thereby allowing quantitative predictions of long-term self-organisation in individual animals. This enables the synthesis of artificial agents that explicitly recapitulate brain-like learning. By doing so, we provide a formal demonstration of the predictive power of active inference, extending beyond descriptive accounts to predict changes in the internal hypotheses underlying individual behaviour. Having said this, the present work is limited to neural data recorded from the telencephalon. Future studies incorporating activity from other brain regions, such as the basal ganglia, may yield more biologically realistic circuit models and improve predictive accuracy.

The reverse engineering framework can be generalised beyond zebrafish and is broadly applicable to diverse animal models, brain regions, tasks, and measurement modalities. The resulting agent can generate theoretical predictions, whose consistency with data serves as a test of the underlying principle. Canonical neural networks can also in principle be extended to hierarchical active inference [44,45], implying potential applications of our method to more complicated tasks. In future work, we plan to apply this framework to mammalian systems and integrate multimodal datasets to investigate whether shared generative structures exist across species, thereby potentially revealing the conserved principles of neural computation.

Moreover, this approach enables a data-driven phenotyping of individual animals based on deviations from Bayes-optimal generative models [46]. Agents dominated by rigid, suboptimal prior beliefs failed to learn the task. As we demonstrated, such individual differences in learning outcomes could be traced back to the initial circuit structure. This enables us to capture the intrinsic mechanistic distinctions between learners and non-learners and the likelihood of task acquisition only from the initial network properties. This may have profound implications for our understanding of how initial conditions and subjective risks mediate the plasticity of neural circuits. Combining mathematical analysis and empirical observation indicates that prior beliefs encoded at the circuit level such as initial synaptic weights are crucial for learning. The notion that differences in prior beliefs can predict subsequent inference and learning may provide insight into the mechanisms underlying inference attenuations and learning disabilities caused by psychiatric disorders [5,47].

In summary, we demonstrated that reverse engineering generative models from neural activity data enables long-term prediction of learning and neuronal self-organisation in zebrafish. These observations provide empirical support for the free-energy principle and active inference as predictive theories of brain function. More broadly, these outcomes highlight the potential of reverse engineering to forecast flexible, individual-specific responses, even in untrained scenarios, which is a key requirement for translational applications in neuroscience and neurotechnology. Our approach offers a powerful tool for uncovering the computational principles of brain function, with potential implications for predicting progression of neurodegenerative and psychiatric disorders and efficiency of education, and for the design of neuromorphic artificial interference that can learn and adapt as the biological brain does.

## METHODS

### Data curation

In this work, previously published data of zebrafish engaging in a go/no-go decision-making task [33] were analysed using the reverse engineering method described in the subsequent sections. The zebrafish were head-fixed under a two-photon microscope for calcium imaging and surrounded by monitors on their front, left, right, and bottom sides, creating a virtual reality environment. Tail movement was tracked using a camera, which allowed the fish to move forward in the virtual space in response to the detected motion. To indicate forward movement, black stripes were displayed on the coloured background, which moved backwards as the fish advanced. A trial continued 10 s, followed by a 15-s rest period. Further details on the preparation of animals, virtual reality settings, two-photon calcium imaging, data acquisition, and preprocessing of data are provided in the previous paper [33].

### Data preprocessing

For analysis of neural activity data measured using two-photon calcium imaging, regions of interest (ROI) were detected as described previously [33]. The dF/F signals after removing low-frequency components using a digital high-pass filter were used for subsequent analyses. Neural activity with this preprocessing is denoted as 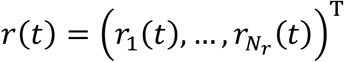, where *N*_*r*_ denotes the number of neurons.

For behavioural analysis, the distance between the start and goal positions was defined as one unit, and the position of the fish was limited in the range of 0 and 2.

### Covariance analysis

Covariance analysis was conducted to investigate the amount of external information encoded by the measured neural activity. To this end, a deconvolutional filter [48] was applied to the ROI data to remove background noise and obtain the estimated spike time-series.

In this setting, the environmental system was characterised by nine state variables: go&blue, go&red, no-go&red, no-go&blue, interval (white), optic flow, electrical stimuli, actions, and position. We assumed that neural activity could be expressed as a weighted sum of these nine state variables and one constant. Some of these variables were not inherently encoded in the circuit initially, but were encoded subsequently through learning, and therefore, their amplitudes changed during the task. Hence, by using a discrete cosine transform (DCT), we considered basis functions that represent changes during a task. Here, a low-dimensional DCT with up to a fifth-order basis was considered, yielding a total of 60 dimensional bases.

Further, we employed a general linear model and optimised the transformation matrix by minimising the mean squared prediction error with least absolute shrinkage and selection operator (LASSO) regression. Then, the amplitudes of the neural activity variances explained by external information were computed. The residuals were treated as noise.

### Statistical tests

The two-sided Wilcoxon signed-rank test was used for the paired comparisons. The two-sided Mann–Whitney *U* test was used for unpaired comparisons.

### Canonical neural networks

We modelled zebrafish neuronal networks (i.e., synthetic agents) using canonical neural networks of rate-coding models comprising two middle-layer and one output-layer neurons [15– 18] (**Fig. 2b**, left). The middle layer neurons *x*(*t*) = (*x*_1_ (*t*), *x*_2_ (*t*))^T^ involve a recurrent circuit, and the output layer neuron *y*(*t*) forms a feedforward network that receives inputs from the middle layer to generate a feedback response to the external milieu. Elements of *x*(*t*) and *y*(*t*) take values within the range of 0 and 1.

Synthetic fish agents received four-dimensional sensory inputs: blue, red, and flow visual inputs, and electric stimuli, expressed as *o*(*t*) = (*o*_*blue*_(*t*), *o*_*red*_(*t*), *o*_*flow*_(*t*), *o*_*stim*_(*t*))^T^. Upon receiving *o*(*t*), the neural activity is expressed as follows [16]:

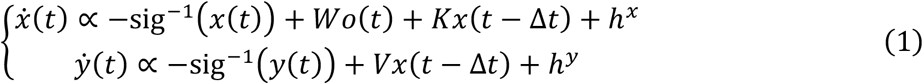

where sig^−1^(·) denotes the leak current in the form of the inverse sigmoid (a.k.a., logit) function; *W* ≔ *W*_1_ *− W*_O_ ∈ ℝ^2×4^, *K* ≔ *K*_1_ *− K*_O_ ∈ ℝ^2×2^, and *V* ≔ *V*_1_ *− V*_O_ ∈ ℝ^1×2^ are synaptic weight matrices for feedforward and recurrent connections in the middle layer and feedforward connections in the output layer, respectively; 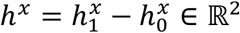 and 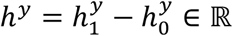 are adaptive firing thresholds in the middle and output layers, respectively; and Δ*t* > 0 is a delay in signal propagation. This model is biologically plausible, as it can be derived from the Hodgkin− Huxley equations [49] and FitzHugh−Nagumo model [50,51] through some approximations [16]. The *W*_1_, *K*_1_, *V*_1_ and *W*_O_, *K*_O_, *V*_O_ matrices can be associated with excitatory and inhibitory synapses, respectively. The adaptive firing thresholds satisfy 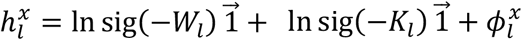 and 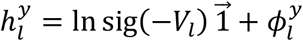. For simplicity, sets of synaptic weights and firing thresholds are defined as *ω* = {*W*_1_, *W*_O_, *K*_1_, *K*_O_, *V*_1_, *V*_O_} and 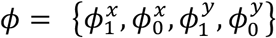, respectively. Throughout the manuscript, ϕ is fixed over the training period, whereas *ω* exhibits synaptic plasticity for each trial.

Without loss of generality, equation (1) can be derived as a gradient descent on a Helmholtz energy (**Fig. 2c**, top left). Following the treatment in the previous work [15,16], the Helmholtz energy for canonical neural networks can be reverse engineered by computing the integral of the right-hand side of equation (1) as follows:

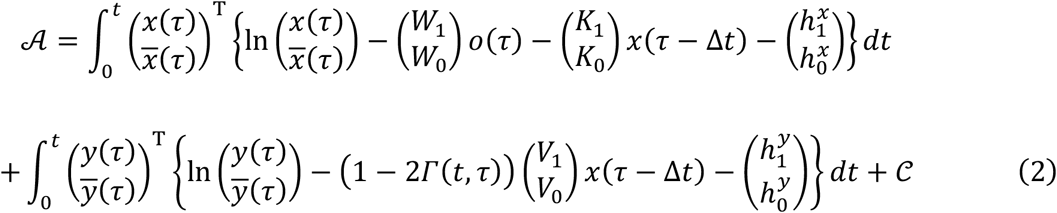

where 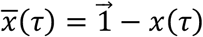 denotes the sign-flipped *x*(τ) centred on 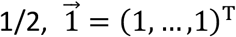 is a vector of ones, and the integration constant 𝒞 is in the order less than *t*. Risk-encoding neuromodulator *Γ*(*t*, τ) that takes the value in the range of 0−1 was added to incorporate the effect of aversive neuromodulations on synaptic plasticity [23−25]. In this work, the risk was determined by the ON or OFF status of punishment (i.e., weak electric stimuli): *Γ*(*t*, τ) = 0 for successful trials in which fish avoids punishment or τ is sufficiently close to *t*; otherwise, *Γ*(*t*, τ) = *Γ*_1_ > 0. The value of *Γ*_1_ varies with individuals. For simplicity, we denote *Γ*_1_ simply as *Γ* in the remainder of the manuscript, except where explicitly stated otherwise. Based on equation (2), equation (1) can be expressed as the gradient descent on 𝒜, i.e., 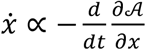 and 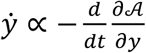.

Then, the synaptic plasticity rules can be derived as a gradient descent on 𝒜 (**Fig. 2c**, bottom left). By computing the derivatives of 𝒜 with respect to synaptic connections, we obtain the following equations for conjugate synaptic plasticity:

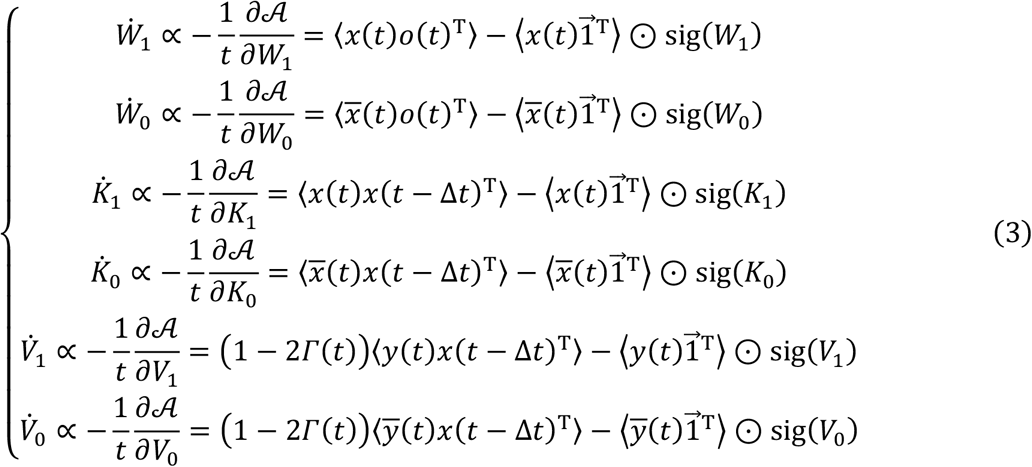

where ⊙ denotes the elementwise (i.e., Hadamard) product operator, 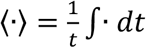 indicates the average over time, and *Γ*(*t*) = ⟨*Γ*(*t, τ*)⟩ denotes the risk for each trial (*Γ*(*t*) = 0 for successful trials and *Γ*(*t*) = *Γ* for failed trials). In detail, by incorporating the initial synaptic weights 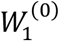 implicit in the integral constant 𝒞, the expectations in equation (3) are computed as 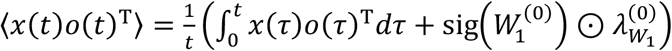 and 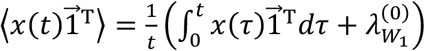 where 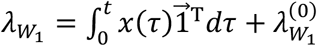 denotes the insensitivity to plasticity (inverse learning rate) and 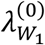 indicates its initial value [15]. The forms of equation (3) are biologically plausible as they comprise Hebbian and homeostatic plasticity, which indicates the biological plausibility of 𝒜 as the energy function that governs canonical neural networks. Synaptic plasticity in *V*_1_ and *V*_O_ is altered by a modulatory factor that encodes the risk *Γ*(*t*), which enables canonical neural networks to optimise actions *δ*(*t*) for avoiding punishment [16]. Further details are provided in previous works [15,16].

### Variational Bayesian inference

Variational Bayesian inference [12] is a process that updates prior beliefs *P*(*ϑ*) about external milieu states *ϑ* to the corresponding approximate posteriors *Q*(ϑ), based on a sequence of observations *o*_1:*t*_ = {*o*_1_, …, *o*_*t*_} (**Fig. 2b**, right). This inference rests upon a generative model expressed as *P*(*o*_1:*t*_, *ϑ*) = *P*(*o*_1:*t*_|*ϑ*)*P*(*ϑ*). Here, a partially observable Markov decision process (POMDP) is employed as a generative model to express a discrete space−time environment [52− 54].

Observations *o*_*τ*_ are generated from hidden states *s*_*τ*_ through likelihood matrix *A* in terms of a categorical distribution *P*(*o*_*τ*_|*s*_*τ*_, *A*) = Cat(*As*_*τ*_) (**Fig. 2a**). The dynamics of hidden states *s*_*τ*_ are determined by transition matrix *B* as *P*(*s*_*τ*_|*s*_*τ*−1_, *B*) = Cat(*Bs*_*τ*−1_). On receiving observations, the agent infers external states and generates actions *δ*_*τ*_ following policy matrix *C* as *P*(*δ*_*τ*_|*s*_*τ*−1_, *C*) = Cat(*Cs*_*τ*−1_). The evaluation of past decisions are conducted based on 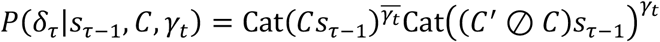using binarized risk *γ*_*t*_ ∈ {0,1}; that is, *P*(*δ*_*τ*_|*s*_*τ*−1_, *C, γ*_*t*_) = Cat(*Cs*_*τ*−1_) for *γ*_*t*_ = 0 and *P*(*δ*_*τ*_|*s*_*τ*−1_, *C, γ*_*t*_) = Cat((*C*^′^ ⊘ *C*)*s*_*τ*−1_) for *γ*_*t*_ = 1, where ⊘ denotes the element-wise division and *C*^′^ is a normalisation factor [16]. The binarized risk *γ* is sampled from a categorical distribution 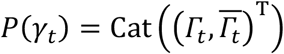; thus, *Γ*_*t*_ is viewed as the risk intensity. Here, *s*_*τ*_ and *δ*_*τ*_ take binary (0,1) values, and parameters *A, B*, and *C* take continuous values between 0 and 1. Thus, the generative model is expressed as

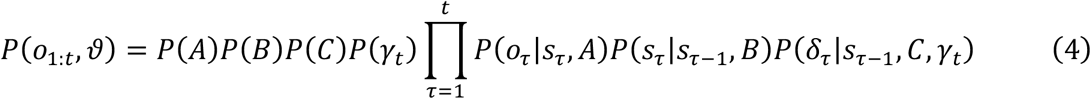

where *ϑ* = {*s*_1:*t*_, *δ*_1:*t*_, *A, B, C*} denotes a set of latent (external) variables.

Variational Bayesian inference minimises variational free energy ℱ[*Q*(*ϑ*), *o*_1:*t*_] = E_*Q*(*ϑ*)_[− ln *P*(*o*_1:*t*_, *ϑ*) + ln *Q*(*ϑ*)] defined as a functional of approximate posterior belief *Q*(*ϑ*) as a tractable proxy for minimising surprise − ln *P*(*o*_1:*t*_) of sensory inputs, where *P*(*o*_1:*t*_) = ∫ *P*(*o*_1:*t*_| *ϑ*)*P*(*ϑ*)*dϑ* denotes the model evidence [13,14]. In POMDP, the approximate posterior belief is given as

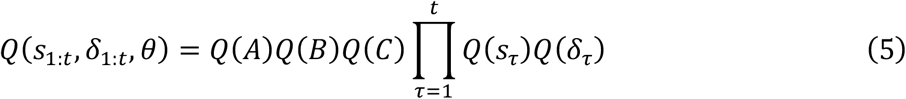

where *Q*(*s*_*τ*_) = Cat(**s**_*τ*_) and *Q*(*δ*_*τ*_) = Cat(**δ**_*τ*_) are categorical distributions and *Q*(*A*) = Dir(**a**), *Q*(*B*) = Dir(**b**), and *Q*(*C*) = Dir(**c**) are Dirichlet distributions parameterised by concentration parameters (Dirichlet counts) **a, b**, and **c**. Based on them, *Q*(*ϑ*) is characterised by the posterior expectation or its counterpart **ϑ** = {**s**_1:*t*_, **δ**_1:*t*_, **a, b, c**}. Hence, the variational free energy ℱ can be analytically expressed as a function of **ϑ** as follows:

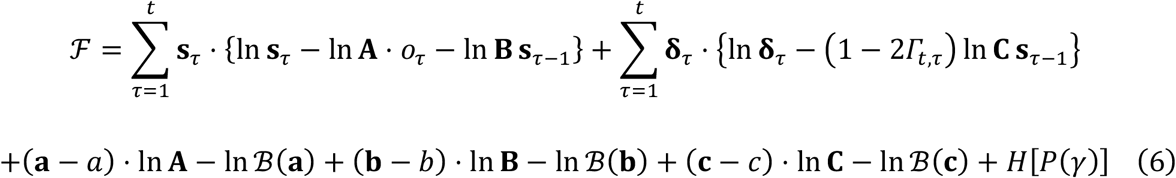

where *P*(*A*) = Dir(*a*), *P*(*B*) = Dir(*b*), and *P*(*C*) = Dir(*c*) are prior Dirichlet distributions, 𝒟_KL_[*Q*(*A*)||*P*(*A*)] = ln **A ⋅** (**a** *− a*) − ℬ(**a**) up to a constant, ℬ(**a**) denotes the beta function, and *H*[*P*(*γ*)] is the entropy of the risk [16]. Here, *Γ*_*t,τ*_ = 0 for successful trials or *τ* = *t*; otherwise, *Γ*_*τ,t*_ = *Γ*.

Minimising ℱ yields an approximate posterior belief *Q*(*ϑ*) that approximates the solution of Bayes’ theorem (**Fig. 2c**, right). Implicit variational update rules are presented as a derivative of ℱ with respect to **ϑ**, and their fixed points provide the posterior expectations. By solving the fixed points *∂*ℱ/*∂***s**_*t*_ = 0 and *∂*ℱ/*∂***δ**_*t*_ = 0, the posterior expectations about hidden states and actions are obtained as follows (**Fig. 2c**, top right):

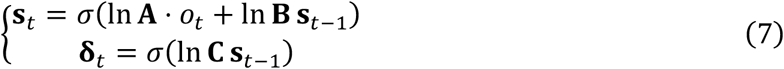

Minimisation of ℱ with respect to parameters, by solving *∂*ℱ/*∂***a** = *O, ∂*ℱ/*∂***b** = *O*, and *∂*ℱ/*∂***c** = *O*, entails the following update rules (**Fig. 2c**, bottom right):

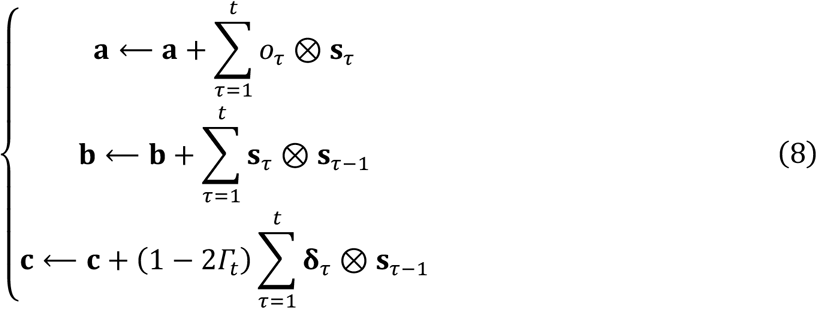

where ⊗ denotes the Kronecker product operator. Parameter posteriors are computed as ln **A**_·*l*_ = ψ(**a**_·*l*_) − ψ(**a**_1*l*_ + **a**_O*l*_) ≈ ln(**a**_·*l*_ ⊘ (**a**_1*l*_ + **a**_O*l*_)) using the digamma function ψ(·). An analogous calculation is used for the other parameter posteriors, **B** and **C**.

Previous work established a mathematical equivalence between the class of factorial POMDP considered here and canonical neural networks (equations (1) and (3)) by showing a one-to-one correspondence between the components of the network’s Helmholtz energy (equation (2)) and the variational free energy (equation (6)) [15,16]; that is, 𝒜 ≡ ℱ, where internal states *φ* encodes posterior beliefs **ϑ** (**Fig. 2c**). For instance, neural activity *x*(*t*) corresponds to the state priors 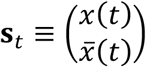 and synaptic weights *W*_1_ and plasticity insensitivities 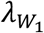 are linked with the parameter posteriors as **A**_1_ ≡ sig(*W*_1_) and 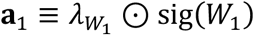 (refer to previous work [15,16] for details). **Table 1** summarises these correspondences. Because all components of canonical neural networks can be mapped into those in variational Bayesian inference under POMDPs, one can reverse engineer the generative models (i.e., internal hypotheses about external milieu) under which fish operate. Through lens of this equivalence, their neural activity and synaptic plasticity are read as performing active inference by minimising variational free energy [15–18].

In particular, delayed modification of synaptic plasticity [19–22] in the form of a three-factor learning rule [23–25] forms optimal behaviour that minimises future risks through the post-hoc evaluation of past decisions. The three-factor learning mediates switching of Hebbian and anti-Hebbian plasticity, which facilitates active inference to acquire risk-minimising behaviour [16].

### Reverse engineering of generative models

This section elaborates an extension of reverse engineering method [26,27] for identifying the generative models of behaving animals (**Fig. 2d**). Crucially, the present scheme estimates the values of the unknown variables—including neural ensemble activity *x*(*t*), effective synaptic weights *W, K, V*, firing threshold factors ϕ^*x*^, ϕ^*y*^, and subjective risk *Γ*—by minimising the shared Helmholtz energy 𝒜 defined in equation (2). Because 𝒜 can be read as the variational free energy, such estimates are identical to posterior expectations of the variational Bayesian inference [26,27], enabling the extraction of a low-dimensional sub-space of ensemble activity *x*(*t*) that maximises the predictability.

Here, the dF/F signals *r*(*t*), sensory inputs *o*(*t*), and actions *δ*(*t*) are the empirically observable variables. A set of these data are denoted as

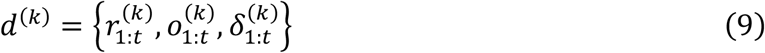

for trial *k*, where *r*_1:*t*_ = {*r*(1), *r*(2), …, *r*(*t*)} expresses a sequence of neural activities (i.e., df/f signals) recorded with a time resolution of 0.1 s. The output layer activity *y*(*t*) is supposed to generate the tail movement of fish *δ*(*t*) through a categorical distribution 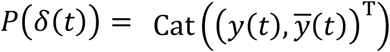. As *y*(*t*) represents the magnitude of the tail movement, *y*(*t*) = *δ*(*t*) is substituted in the following analysis.

Empirical neural activity data are assigned to canonical neural network models, and the corresponding generative models are identified through the established equivalence. This process is begun by classifying large-scale neural activity data into two-dimensional neuronal ensembles. The neuronal ensemble activity *x*(*t*) is characterised by the mapping from individual neural activity to ensemble activity, expressed as 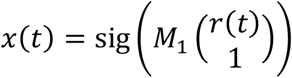 with the values in the range between 0 and 1. A reconstruction mapping from *x*(*t*) to individual neural activity is also defined as 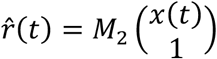, where *M* = {*M*_1_, *M*_2_} denotes encoding and decoding matrices for classification. Moreover, the characterisation of the network architecture requires to specify unobservable network parameters such as synaptic weights (*ω*) and firing thresholds (ϕ). Thus, the internal states of canonical neural networks for trial *k* are provided as follows:

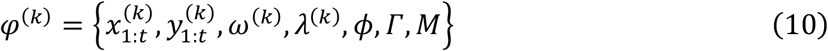

where 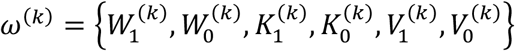 denotes effective synaptic weights, 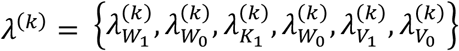 denotes plasticity insensitivity (stability of synaptic weights), 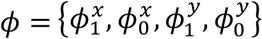 denotes firing threshold factors, and *Γ* is the subjective risk for receiving weak electric stimuli. Here, ϕ, *Γ*, and *M* are fixed over trials. The magnitude of plasticity is controlled by plasticity insensitivity *λ*.

The allocation into neuronal ensembles makes the neural activity suitable for analysis. This is conducted by optimising classification matrices *M* = {*M*_1_, *M*_2_} to minimise 𝒜, following the gradient descent:

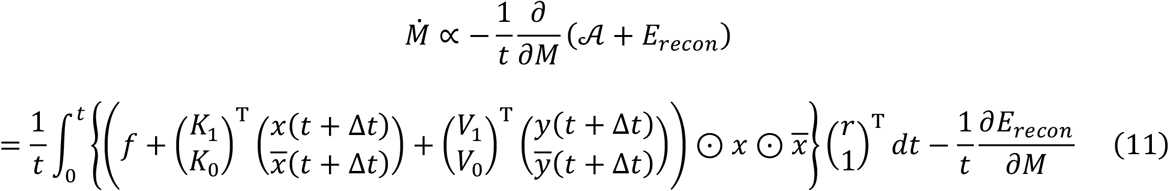

where *f* = −sig^−1^(*x*(*t*)) + *Wo*(*t*) + *Kx*(*t −* Δ*t*) + *h*^*x*^ is the right-hand side of equation (1). To preclude pseudo-solutions, a reconstruction error term 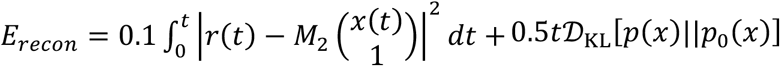 was added to ensure the efficient extraction of major features in neural activity data. The Kullback−Leibler divergence with an independent uniform prior distribution *p*_O_(*x*) prevents ensembles *x* from mutually correlating too strongly.

Projecting empirical neural data onto the manifold of *x* enables interpretable, mechanistically grounded predictions of learning trajectories. When estimating the generative model (**Fig. 2d** and **Fig. 3**), the allocation was conducted using data in all 100 trials, *d*^(1)^, …, *d*^(1OO)^. Conversely, when predicting learning trajectories (**Fig. 2e** and **Fig. 4**), this was conducted using test data in the initial 20-trial adaptation period, *d*^(1)^, …, *d*^(2O)^, to avoid double-dipping.

For the firing threshold factors, we considered that ϕ^*x*^ and ϕ^*y*^ were constant during the short experimental period and derived their posterior expectations as follows:

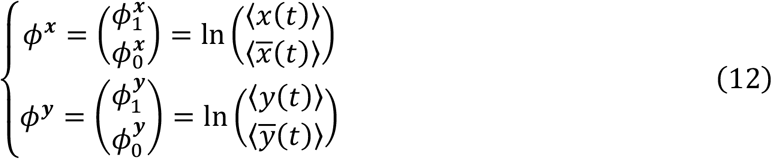

where ⟨·⟩ denotes the average (i.e., the mean value) over the data during the initial 20-trial adaptation period.

Effective synaptic connectivities (*W, K, V*) were altered through activity dependent plasticity.

Thus, these weights were estimated for each trial by minimising 𝒜. Solving the fixed point of equation (3) entails the following synaptic weights that minimise 𝒜:

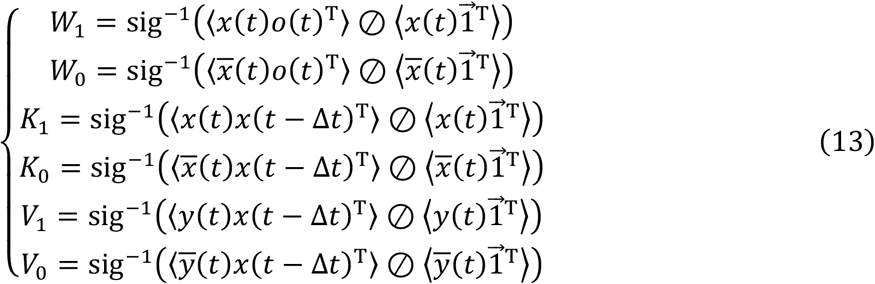

where ⊘ denotes the element-wise division and ⟨·⟩ indicates the average over data sequences up to the current trial. Substituting empirical data sequence up to trial *k, d*^(1)^, …, *d*^(*k*)^, into equation (13) yields the effective synaptic connectivity for trial *k*.

The subjective risk was estimated using Bayesian model selection. Here, the variational free energy ℱ (equivalent to 𝒜) was computed with varying *Γ* for receiving aversive electric stimuli between 0 and 1, and the value of *Γ* that minimise ℱ over training period was selected.

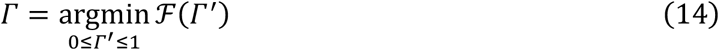

This enables the characterisation of individual differences in punishment-related learning.

In summary, the initial network states *φ*^(O)^ = {*ω*^(O)^, *λ*^(O)^, ϕ, *Γ, M*} at trial 0 comprise the classification matrix *M*, initial synaptic weights *ω*^(O)^, insensitivity to plasticity *λ*^(O)^, firing threshold factors ϕ, and subjective risk *Γ*. Firing threshold factors ϕ, initial synaptic weights *ω*^(O)^, and initial plasticity insensitivity *λ*^(O)^ were determined as values that minimise 𝒜 by using the empirical data in the initial 20 trials, *d*^(1)^, …, *d*^(2O)^, whereas *Γ* and *M* were determined using data during the entire training period, *d*^(1)^, …, *d*^(1OO)^. These variables are computed by executing the naive initialisation function that involves equations (11)−(14) (refer to reverse_engineering.m and phi_init.m in Code [55]):

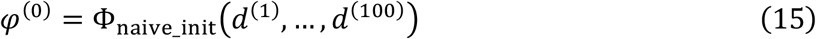

Moreover, the estimation of internal variables in trial *k* (1 ≤ *k* ≤ 100) was defined as a mapping from *φ*^(*k−*1)^ to *φ*^(*k*)^ given observable data *d*^(*k*)^ and performed by executing the following estimation function that computes equation (13) (see phi_estimate.m in Code [55]):

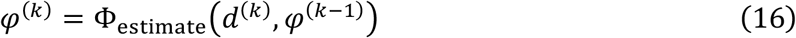

By recursively computing this function, ensemble activity 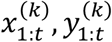, synaptic weights *ω*^(*k*)^, and plasticity insensitivity *λ*^(*k*)^ were updated for each trial, whereas ϕ, *Γ*, and *M* remained fixed throughout the training period. Owing to the established equivalence between canonical neural networks and variational Bayesian inference [15−18], the internal states *φ*^(*k*)^ can be conceptualised in terms of posterior beliefs about external states **ϑ**^(*k*)^ (**Table 1**).

### Prediction of learning trajectories

This section establishes long-term prediction of internal states based on the free-energy principle (**Fig. 2e**). When the free-energy principle is applied to a particular system, its predictive validity can be examined by asking whether it can predict the system response [26]. On this reading, a key challenge lies in forecasting long-term learning and self-organisation at the level of individual brains. Such predictions are particularly challenging under closed-loop conditions considered herein, because external inputs change depending on the actions of the fish, and this generally yields largely different data sequences. The reverse engineered generative model can help us overcome this by integrating bottom-up statistical inferences with theoretical predictions.

The predicted neural activities 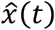 and ŷ (*t*) were provided by substituting predicted variables (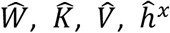, and 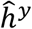) into equation (1), as follows:

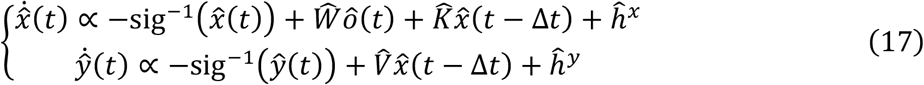

where 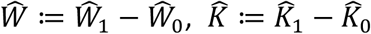, and 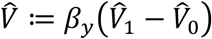 denote predicted synaptic weight matrices, 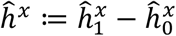 and 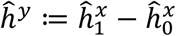 denote predicted firing thresholds that satisfy 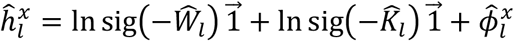 and 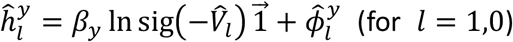, and *β*_*y*_ = 3 represents a gain parameter introduced to align predicted activity more closely with empirical activity. Here, unlike parameter estimation in the previous section, both the neural activity and synaptic weights are determined to minimise 𝒜 in the absence of empirical data. Moreover, owing to the closed-loop condition, predicted inputs ô(*t*) differ from empirical data *o*(*t*) as the agent’s action changes the environmental states. Given these, the updates of synaptic strengths were determined as follows:

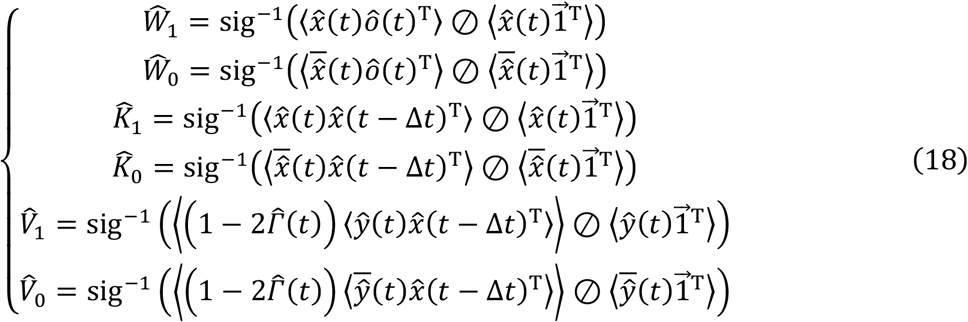

Using equations (17) and (18), the predicted neural activity and synaptic strengths were computed for each trial to obtain the sequences of neural activity and synaptic strengths.

To achieve accurate predictions, precise characterisation of the initial network states is crucial. However, this is not straightforward because equation (15) relies on the empirical data from the entire training period; thus, simply applying equation (15) to predictions would constitute double dipping. To overcome this, we developed a pretrained initialisation function that predicts the initial parameters of canonical neural networks based only on the initial empirical data, which is defined as follows (phi_pretrained_init.m in Code [55]):

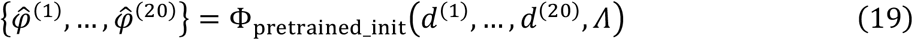

where *Λ* denotes a set of hyper-parameters. Here, predicted initial states 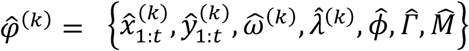 include neural activity 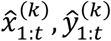, classification matrix 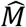, initial synaptic weights 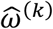, initial plasticity insensitivity 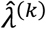, firing thresholds 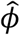, and subjective risk 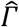. This function outputs the predicted initial states 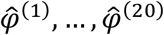 based on observable data in the initial 20-trial adaptation period, *d*^(1)^, …, *d*^(2O)^. First, 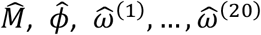, and 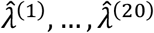 were estimated using equations (11)−(13) based on the initial data *d*^(1)^, …, *d*^(2O)^ of a fish. Then, 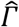 was estimated and 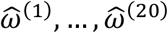 and 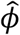 were modified using a function 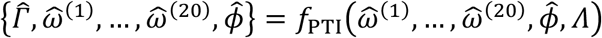.

Hyper parameters *Λ* were optimised to maximise the predictability of the states up to trial 100 using the training dataset, which was conducted by minimising errors between empirical and predicted variables over training dataset. The prediction analysis throughout this paper was conducted based on leave-one-out cross-validation method [40]. To avoid double-dipping data when making predictions, *Λ* was trained using data from trials 1 to 100 in the 44-fish training dataset. Then, the predicted initial values 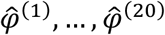 for the test data were estimated using only data from the initial 20 trials of the test fish.

Trained on an activity dataset of other fish, this function estimates the initial internal variables for each fish based on the neural activity during the initial adaptation phase. Hence, while a part of parameters such as subjective risk *Γ* cannot be computed directly from the initial observable data as fish had not yet received punishment, the pretrained initialisation allow to characterise the initial states of the test data. This method automates the identification of neural ensembles and the selection of initial parameter values, which were previously performed heuristically [27], and improves model initialisation.

After the initial states 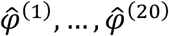 are determined, the canonical neural network model simulates the subsequent activity and plasticity according to the free energy minimisation, which yields a series of internal states 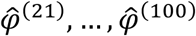 and external states 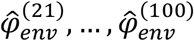. These predictions were performed by executing the prediction function that involves equations (17) and (18) (see phi_predict.m in Code [55]):

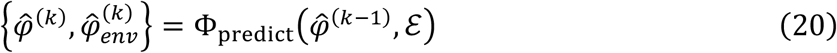

where ℰ = {*A, B, C, D, E*} denotes a set of parameters that determine the POMDP environmental settings and 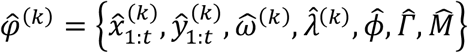 and 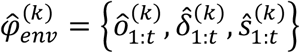 are predicted internal and external states at trial *k*, respectively. These predicted internal states 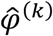 were then compared with the sequences of empirical internal states *φ*^(*k*)^ to assess the prediction accuracy (**Fig. 4**).

## Data Availability

Example raw data analysed in this work are available in the repository (https://doi.org/10.5281/zenodo.5195611). Full raw data are available from the corresponding author of previous work [33] upon request. Source data are provided with this paper.

## Code Availability

The simulations and analyses were conducted using MATLAB version R2020a. The scripts are available at GitHub https://github.com/takuyaisomura/reverse_engineering [55]. The scripts are covered under the GNU General Public License v3.0.

## Acknowledgements

T.I. is supported by the Japan Society for the Promotion of Science (JSPS) KAKENHI under Grant Number JP23H04973, the Japan Agency for Medical Research and Development (AMED) under Grant Number JP23wm0625001, and the Japan Science and Technology Agency (JST) CREST under Grant Number JPMJCR22P1. The funders had no role in study design, data collection and analysis, decision to publish, or preparation of the manuscript.

